# Genotype-level quality control substantially reduces error rates in population-scale whole-genome sequencing

**DOI:** 10.1101/2025.11.06.686913

**Authors:** V. Kartik Chundru, Harrison I. W. Wright, Timothy M. Frayling, Andrew R. Wood, Michael N. Weedon, Caroline F. Wright, Robin N. Beaumont, Gareth Hawkes

## Abstract

Population-scale whole-genome sequencing data will contain many individual-level genotype errors, even after allele-level quality control (QC). We establish the need for genotype-level QC using UK Biobank (N=490,726) and All of Us v8 (N=414,830), where we remove up to 100 million (∼9%) additional low-quality variants. We demonstrate reduced false positive rate in downstream genetic association studies, highlight the power of parent-offspring trios for QC, and illustrate the need for sex-specific X-chromosome filtering. We provide a QCed All of Us v8 dataset in *plink*-pgen format, and an efficient pipeline for QC and conversion from VCF to *plink-*pgen for UK Biobank.

## Main

Between 2018 and 2025, over one million high-coverage short-read whole-genome sequences (WGS) have been released publicly across population-scale biobanks such as UK Biobank (UKBB)^1^, All of Us v8 (AoUv8)^2^ and TOPMed^3^. Variant-level quality control (QC) for sequencing data has recently evolved to be applied at an allele-level using machine learning approaches such as GATK VETS (Variant Extract Train Score)^4,5^, the successor to VQSR (Variant Quality Score Recalibration), and the DRAGEN ML-score developed for the UKBB WGS release^6^. These methods classify each alternate allele, which after multi-allelic splitting are each interpreted as a single variant, as PASS/FAIL which allows for the situation where a multiallelic variant has alternate alleles of mixed qualities. However, an issue overlooked by these allele-specific methods is where the *average* quality of an allele/variant is high, but there are a number of individuals with low-quality genotype calls (examples given in **Supplementary Figure 1**). This is particularly relevant for ultra-rare variants identified by population-scale WGS studies, where most genotypes are homozygous-reference and easier to call, which could substantially impact rare-variant burden testing (e.g. **Supplementary Figure 2**). While some studies have discussed the benefits of applying these filters^7^, there are no standard strategies to evaluate their effectiveness and establish whether optimal thresholds differ across cohorts, and no cost effective way to apply them to population-scale datasets where they are not applied by default (e.g. UKBB and AoUv8).

Here, we present a framework for implementing genotype-level QC and assess the error rates across QC approaches in population-scale WGS data using UKBB and AoUv8, across both the autosomes and X-chromosome (see **Figure 1** for study design). We release an efficient pipeline to QC and convert the UKBB sharded pVCFs and AoUv8 VDS data to plink-*pgen* format, the latter of which we provide in a public workspace for registered users.

**Figure 1.**
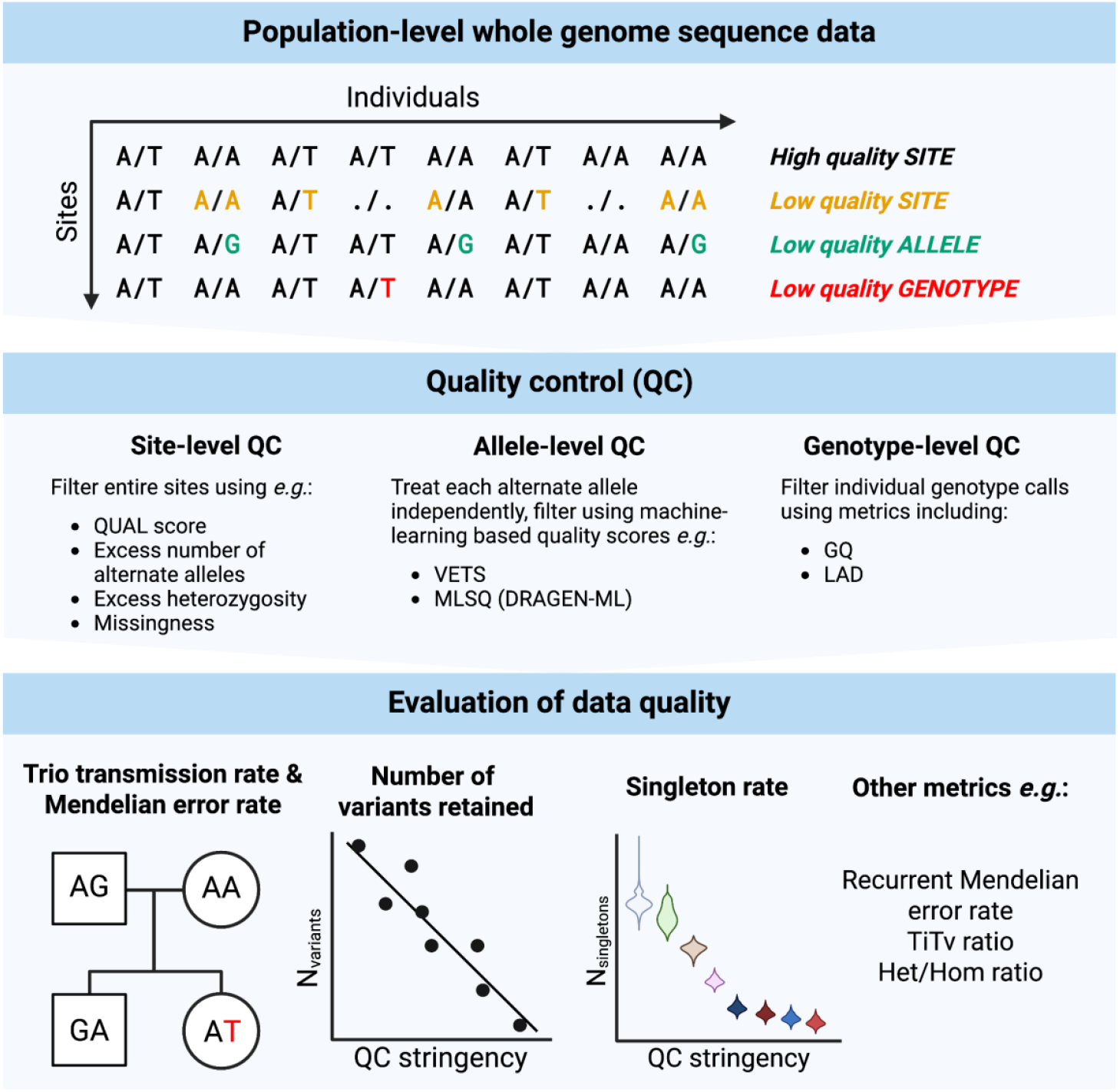
Study Design. A pictorial representation of the study design.

We compared the provided allele-level filtered UKBB (N=490,276; UKBB-DRAGEN-ML), and AoUv8 (N = 414,830; AoU-VETS) WGS data against a second version of these datasets with additional genotype-level QC (UKBB-GenotypeQC and AoU-GenotypeQC). We selected a set of genotype-level QC thresholds by varying the genotype quality (GQ) and sequence depth (DP) at which a genotype was set to missing, and subsequent filters on the allowed genotype missingness rate, on 100 randomly-selected autosomal pVCF chunks of ∼20kb each (**Methods**; **Supplementary Figure 3**). For the X chromosome, we used 10 randomly-selected pVCF chunks from the non-pseudoautosomal region, and another 10 across the pseudoautosomal regions (**Supplementary Figure 4**).

Firstly, it is important to note that the GQ value provided by DRAGEN^8^ is different from the GQ exported from GATK^9^, which means thresholds developed for one platform are not applicable to the other. GQ value distributions can also be rescaled differently depending on the downstream joint-called pipeline used (**Supplementary Figure 5**). Secondly, in AoUv8 GQ<20 homozygote reference genotypes were pre-filtered before release, unlike UKBB. Taking these issues into account, with the additional aim of achieving equitable missingness between the two biobanks, we settled on the following thresholds; UKBB: GQ>10, DP>7, and missingness<0.1; AoU: GQ≥20 (in harmony with the existing hom-ref filtering), DP>7, and missingness<0.1.

Using a modification of *plink2*^10^ to calculate total depth from the local allele depth, we converted the whole genome from pVCF shards to *plink*-pgen while also applying genotype-level filtering. Our implementation of genotype-level QC is faster and cheaper compared to conventional approaches using *bcftools*^11^ (∼80% reduced cost, cpu time, and saves a total of 1.75 MWh of carbon per conversion of UKBB WGS data, based on the green-algorithms application^12^, which is equivalent to 37.04 tree-years, i.e. it would take 37 years for an average mature tree to absorb the equivalent carbon).

A total of 1,104,132,700 (91.5%) variants were retained after applying our genotype-level QC thresholds in the UKBB-GenotypeQC set, compared to 1,207,316,256 variants left after only removing variants classified as ‘FAIL’ in the UKBB-DRAGEN-ML set. Overall, ∼0.3% of genotypes were removed from variants that pass allele-level QC, with most genotypes removed being homozygote-reference (**Figure 2A**). Of note, in UKBB the majority of errors were hom-ref calls with low quality, likely due to the high prior of hom-refs used by variant callers. In the AoU-GenotypeQC, a total of 1,331,260,959 (99.3%) variants were retained compared to a total of 1,340,702,367 ‘PASS’ variants in the AoU-VETS dataset. We note that the reduced impact of genotype-level filtering in the AoUv8 dataset is due to their pre-filtering of homozygote reference genotypes with GQ<20 and subsequent allele-level missingness filtering.

**Figure 2.**
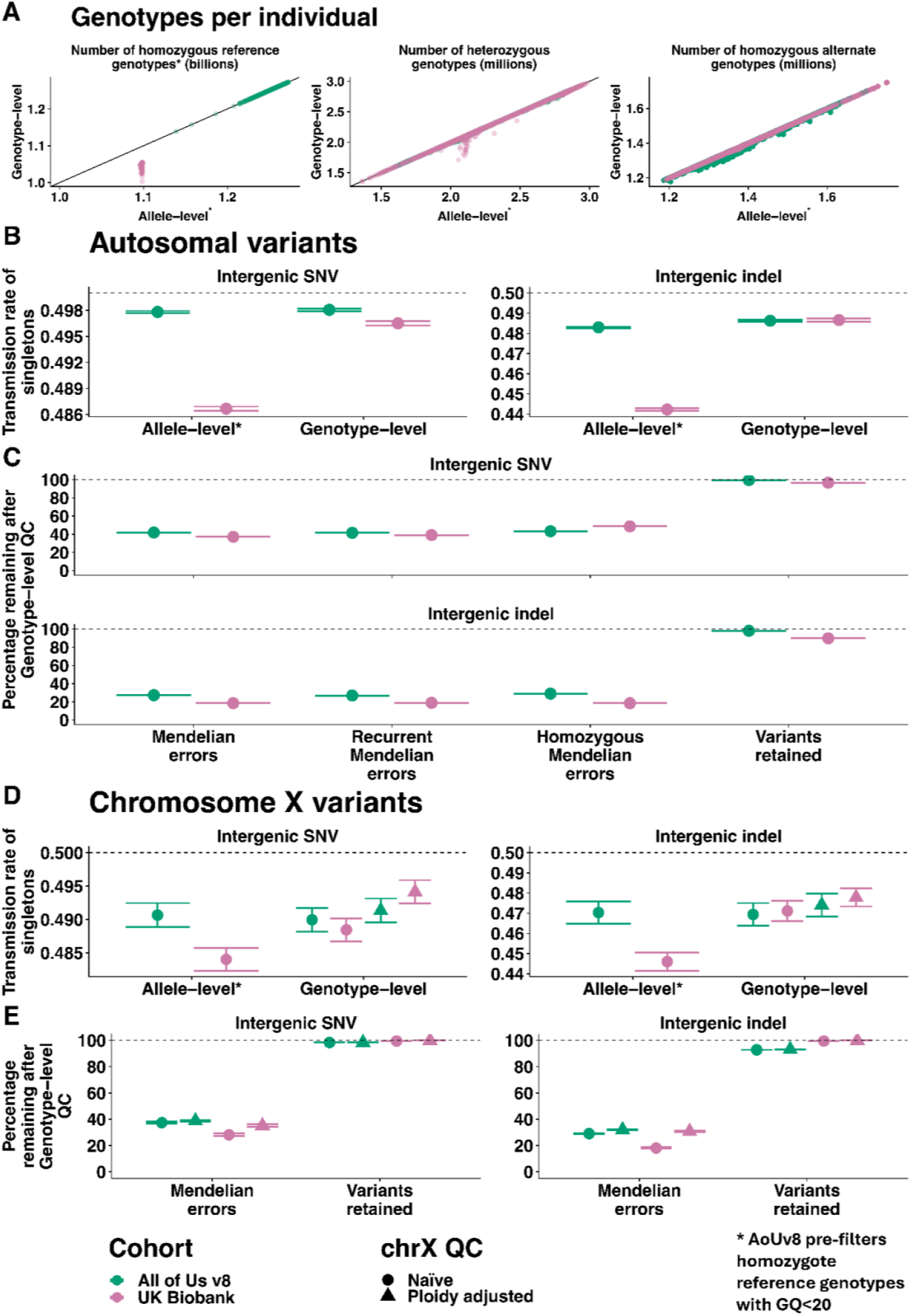
Genotype-level QC improves WGS data quality compared with allele-level QC. A) Comparisons of the number of homozygous-reference, heterozygous, and homozygous-alternative genotypes per individual from the different QC datasets. B) Transmission rate of singleton genotypes in trios for allele- and genotype-level QC datasets of UK Biobanks and All of Us v8. We restrict to intergenic SNVs and indels assuming these are null variants. C) Percentage of errors remaining, and variants retained after genotype-level QC. D) and E) Transmission rate and percentage of errors and variants retained for male proband trios with the shape of point denoting whether the genotype-level QC was adjusted for ploidy.

Measuring the error rate in WGS data is not trivial without a known truth-set of genotypes. However, under the assumption of no selection acting upon intergenic variants, genotypes should be transmitted from parents to offspring at a rate of 50% according to Mendelian inheritance rules. Deviation from this 50% transmission rate would therefore indicate the presence of genotyping errors^13^, making parent-offspring trios in large-scale sequencing data extremely valuable. Restricting to singleton variants only observed within one trio increases sensitivity to observe the differences between filtering thresholds, otherwise common variants would dominate any signal, masking any changes in quality for rare variants. Across all groups of variants, applying genotype-level QC increased the transmission rate closer to the expected 0.5 (**Fig 2B**); an increase of 2.02% and 0.05% for SNVs and 10.02% and 0.68% for indels in UKBB and AoUv8 respectively (**Supplementary Table 1**). In addition to transmission rates, the number of Mendelian errors per trio is a near-direct measure of the number of errors in the dataset given that only a very small proportion of these will be true *de novo* mutations (∼70 per individual^14,15^). The autosomal Mendelian error rate for (presumed null) intergenic variants was substantially reduced after genotype-level QC (**Figure 2B-C**), with a 62.86% and 58.08% reduction for SNVs and 80.42% and 72.66% for indels in UKBB and AoUv8 respectively (**Supplementary Table 1**). Overall, variants in the QC sets were significantly below 0.5 transmission (all p<10^−116^), and Mendelian errors remained, for (presumed null) intergenic variants, even when excluding variants in difficult to sequence regions (**Supplementary Figure 6**). Our results show that despite rigorous QC, the data still contain sequencing errors that are difficult to remove at scale. We therefore recommend manual read-level review^16^ (**Supplementary Figure 1**) when specific or very rare variant calls are integral to a study.

Separate consideration must be given to the X chromosome. In particular, because males are haploid for chromosome X, the sequencing depth will be lower in regions which are not pseudo-observed on chromosome Y (non-Pseudo-Autosomal Regions; non-PAR). To adjust for this we halved the depth threshold for males in the genotype-level QC. We also exclusively considered the transmission of mothers to probands in the non-PAR regions, and focused on trios with male probands to test if adjusting for ploidy was required for QC (**Methods**). When genotype-level QC is applied naïvely, i.e. treating males the same as females, the chromosome X non-PAR transmission rate in trios with male probands increases compared to allele-level QC by 0.91% and 5.62% in UKBB for SNVs and indels respectively, and decreases by 0.15% and 0.19% in AoUv8. When applying genotype-level QC in a sex-specific manner we see that the transmission rate in UKBB is improved by 2.09% and 7.14%, and the transmission rate in AoUv8 now improved by 0.14% and 0.80%, for SNVs and indels respectively (**Figure 2D-E**). With sex-specific QC we retain an additional 0.4% of variants in UKBB, and 0.1% of variants in AoUv8, highlighting the need for sex-specific QC of the X chromosome.

To demonstrate the impact of genotype-QC on downstream analyses, we performed an autosomal single-variant genome-wide-association-study of sex in UKBB, under the hypothesis that variants associated with sex are enriched for poorly-called genotypes due to shared homology with sex-chromosome sequence^17^. We tested all variants with minor-allele-count ≥ 10 after filtering on Hardy-Weinberg *P* in accordance with Greer *et al*.^18^ in individuals of genetically-inferred European ancestry. In total, we observed independently associated genetic variants on 21/22 chromosomes in the allele-level QC dataset, but only 14/22 in the genotype-level QC dataset. Specifically, in the provided allele-level QC UKBB dataset, we identified 231 variants independently associated (*P* ≤ 2.95e-10; **Methods**) with sex, of which 130 (56%) were rare (minor-allele-frequency < 0.1%). Of these 231 variants, 208 (90%) were present in gnomADv4.1^19^ however, 124 (60%) of these failed quality filters in the gnomAD genomes (which do not contain UKBB samples) (**Supplementary Table 3 and 4**). In contrast, after applying genotype-level QC we observed 153 (66%) independently associated variants, of which 12 (8%) were rare. Of the 153 variants, 151 (99%) were observed in gnomADv4.1, with 29 (19%) of these variants failing quality filters in gnomAD genomes. However, we observed a higher rate of ‘jumpers’ (CoJo joint LOG10*P* -marginal LOG10*P*) > 3) in our genoQC results (126; 82%) than for DRAGENML (52; 23%). We hypothesise this is due to our filtering of variation in a given sequencing read which has the highest homology with chrX or chrY, and retain less homologic variants as they will be of higher quality but imperfectly tag the homology signal. As has been shown previously, joint modelling on imperfectly tagged SNPs can cause jumpers^20^.

Other metrics often used for QC of WGS datasets include transition-to-transversion (Ti/Tv) ratio, and heterozygous-to-homozygous ratio. While these metrics are very useful to capture the average quality of variants, we find that they are not sensitive enough to capture errors on the individual-level, with minimal changes after genotype-level QC (**Supplementary text, Supplementary Figure 7**). Another commonly used metric – the number of singleton variants per individual – showed a marked difference for some individuals in UKBB who had outlying levels of singletons before applying genotype-level QC (**Supplementary Figure 8**).

Our study has a number of limitations. Firstly, to choose the different thresholds of GQ, depth, and missingness, we tested a limited and discrete set of values based on prior knowledge on a limited subset of data. Ideally, we would apply an unbiased method to search efficiently across the parameter space for the best set of quality thresholds. Secondly, we did not assess the effects of different filtering thresholds within contexts beyond association testing, such as rare disease diagnosis and model-training, which may require a different balance of stringency against number of variants retained. We therefore emphasise that our findings offer guidance for individual researchers to apply QC to their own data within a given context, where different requirements for sensitivity and specificity will necessitate different thresholds for genotype filtering. Thirdly, we acknowledge that many correct but low-quality genotype calls will be removed in the process of genotype-level QC. This is an unavoidable consequence of applying imperfect filters at a genome-wide scale. For analyses looking at subsets of variants and individuals, it may still be preferable to manually inspect individual calls.

In conclusion, our recommendation is to apply genotype-level filtering on top of the allele-level filters that are generally applied to WGS datasets. Genotype-level filtering should be calibrated to individual cohorts using, if possible, truth sets such as parent-offspring genotype transmission. Additionally, data from haploid males in the non-pseudoautosomal regions of X chromosome should be given more lenient depth filtering to maximise variant retention and avoid potential sex bias in genotype calls. Applying these procedures will reduce false positive calls and increase power to detect novel associations by enabling harmonisation across cohorts.

## Methods

### UKBB WGS Data

The whole genome sequencing performed for UKBB^6^ had an average coverage of 32.5X using Illumina NovaSeq 6000 sequencing technology. The genome build used for mapping was GRCh38: single variant nucleotide variants and short insertions or deletions (indels) were jointly called using DRAGEN^8^ 3.7.8. The UKBB-DRAGEN-ML release was provided in three formats: pVCF, plink 2.0^10^ pgen and bgen. We retained alleles (variants) classified as ‘PASS’ in the pVCF FILTER entry by the DRAGEN-ML algorithm. The pVCF format of the UKB-DRAGEN-ML release did not retain per-genotype GQ and LAD entries: in order to perform additional genotype QC, we used the initial release of UKBB DRAGEN WGS data, which encompassed 154,430 pVCF files across 490,541 individuals. The application of genotype-level QC we used for this analysis was prior to the development of the conversion pipeline we detail in this paper. We used bcftools^11^ to apply genotype-level filters, split multi-allelic sites, and left-aligned all REF and ALT alleles using a 1000G b38 reference available at https://ftp.1000genomes.ebi.ac.uk/vol1/ftp/technical/reference/GRCh38_reference_genome/ (accessed 30/03/2024). We then assigned all variants a unique ID (CHR:POS:REF:ALT), followed by plink2^10^ to convert each filtered pVCF shard to a plink2 format and concatenate into per-chromosome p(gen/sam/var) files.

Based on a KING^21^ analysis done by UK Biobank, we identified 1,040 UKBB-trios by matching individuals inferred to be parent-offspring relations.

### All of Us v8 WGS Data

The whole genome sequencing performed for AoU had average coverage 37.9x using Illumina NovaSeq 6000 sequencing technology. The genome build used for mapping was GRCh38: single variant nucleotide variants and short indels were called using DRAGEN Joint calling was performed via the Genomic Variant Store^22^. The full AoUv8 WGS data was provided exclusively in Hail VDS format^23^, with VETS^4^ filtering flags and site-level filters as described in the documentation^5^. We exported the Hail VDS to VCF format using Hail v0.2.134-952ae203dbbe.

Although the AoUv8 documentation^5^ describes VETS as “genotype-level” it is allele-level and does not differentiate between the same genotype calls in different individuals. Instead, all genotypes with a poor quality allele are flagged. In addition, the VDS format compresses WGS data by storing homozygote reference calls in reference blocks, with blocked-GQ values of {20, 30, 40}, representing a GQ of 20-29, 30-39 and 40+ respectively. Homozygote reference calls with GQ<20 were set to be missing in the released VDS dataset.

Based on a KING relatedness analysis using array data, which matched the relatedness analysis using WGS data released by AoU, we identified 1,321 AoU-trios.

### Identification of optimal quality filters for genotype-level filtering

We varied the genotype-quality cutoffs for GQ (Genotype Quality) and LAD (Local Allele Depth) at which we assigned a genotype to missing, and varied the missingness threshold at which a variant is excluded at the following thresholds: GQ = {10, 15, 20, 25, 30}; LAD = {8,10, 12} and MISS={0.1, 0.5}. It was not computationally feasible to examine the effect of varying genotype-filters on all of the initial-release pVCFs. Instead, we limited our analysis to 100 randomly selected autosomal pVCFs covering two million basepairs, and ten pVCFs in both the chromosome X PAR and non-PAR regions. For chromosome X specifically, we additionally tested the effects of halving the LAD threshold for males, due to their haploid status.

We then performed a single UKBB genome-wide WGS conversion to plink format for the GQ = 10, LAD = 8, MISSINGNESS = 0.1 filters, in addition to the allele-level filtering using DRAGEN-ML, to test that the generalisability of our chunk estimates were accurate, and compare directly genome-wide data with the allele-level QC in UKBB.

To generate the AoU-GenotypeQC dataset, we separately applied a range of custom genotype-level filtering to the same 100 genomic regions described on top of the AoUv8 VETS filter. We used a modified version of plink2 (**Data & Code Availability**) to apply LAD and GQ filters, and to split multi-allelic variants and left-align allele representations (using the same reference as for UKBB). This updated methodology is more computationally efficient, with reduced carbon footprint than using *bcftools*. We set any alleles to be missing if they had GQ<20 or LAD<8, and dropped variants which subsequently had >10% missingness.

### Mendelian Error and Transmission Ratio

We used the bcftools^11^ (v1.22) trio-stats plugin to calculate the autosomal and pseudo-autosomal mendelian error and singleton transmission rates. For estimating the transmission ratio on the non-pseudoautosomal regions chromosome X, we used a custom script (**Data & Code Availability**), and we used plink2^10^ to detect the Mendelian errors.

### Association Testing with sex

We defined a UKB-sex phenotype for individuals whose self-declared and genetically-inferred sex matched (UKB fields 31 and 22001). To identify single variants associated with sex we performed an association test for all genetic variants with a minor-allele-count of at least 10 using REGENIE^24^ (v4.1) in 1,361 pseudo-linkage-independent chunks^25^ of the 22 autosomes, after filtering on Hardy-Weinberg equilibrium (plink2 --hwe 0.05 0.001). Chunk-wise lead variants were then selected in a conditional-joint analysis using an altered version of GCTA-COJO^20,26^ (with command line options diff-freq = 0.2, COJO-P = 2.95×10-10), with the UK Biobank whole-genome sequencing data as the LD reference panel. To define chromosome-wide independently associated variants, we applied a second round of the altered GCTA-COJO algorithm, considering only those variants which were classified as independent at the chunk-level.

## Data & Code Availability

Individual-level data cannot be shared publicly because of data availability and data return policies of the UK Biobank and All of Us. Data are available from the UK Biobank and All of Us for researchers who meet the criteria for access (http://www.ukbiobank.ac.uk; https://allofus.nih.gov/).

We have made our quality-controlled All of Us v8 whole-genome plink dataset available in a community controlled-tier workspace: [to be added upon acceptance]. The UK Biobank DNA Nexus applet for converting the WGS data from pVCF shards to pgen format, the scripts to extract WGS data from Hail VDS format, and other analysis and plotting scripts are available on github at https://github.com/chundruv/ukbb_pvcf2pgen.git, and the plink modifications at https://github.com/chundruv/plink-ukbb-mod.git.

## Acknowledgements

VKC and CFW supported by the Wellcome Trust Grant 226083/Z/22/Z. HW and MNW are supported by the Medical Research Council grant MR/Y003748/1. ARW is supported by the Academy of Medical Sciences / the Wellcome Trust / the Government Department of Business, Energy and Industrial Strategy / the British Heart Foundation / Diabetes UK Springboard Award [SBF006\1134]. TMF is supported by MRC awards MR/WO14548/1 and MR/T002239/1. GH is supported by the Medical Research Council grant UKRI327. The research utilised data from the UK Biobank resource carried out under UK Biobank application number 103356. UK Biobank protocols were approved by the National Research Ethics Service Committee. The authors would like to acknowledge the use of the University of Exeter High-Performance Computing (HPC) facility in carrying out this work, funded by an MRC Clinical Research Infrastructure award (MRC Grant: MR/M008924/1). This study was supported by the National Institute for Health and Care Research Exeter Biomedical Research Centre, and by the UK Medical Research Council [grant number MR/W014548/1]. The views expressed are those of the authors and not necessarily those of the NIHR or the Department of Health and Social Care. We gratefully acknowledge All of Us participants for their contributions, without whom this research would not have been possible. We also thank the National Institutes of Health’s All of Us Research Program for making available the participant data examined in this study

## Author Contributions

VKC, RNB, GH, CFW, MNW and TMF jointly conceived the study. VKC, HIWW, RNB, GH and ARW generated pipelines, and performed QC and analyses. VKC, RNB, HIWW and GH wrote the manuscript. All authors have read and approved the paper.

