## SupplementaryMaterial for "Genotype-level quality control substantially reduces error rates in population-scale whole-genome sequencing"

### Supplementary Information

#### Supplementary Text

##### *Definition of the genotype quality (GQ) metric*

The GQ value is the Phred-scaled probability that a genotype call is incorrect; the classical GATK[1] GQ is capped at 99 which causes a heavy skew to the maximum value, and most studies remove values at GQ<20. In DRAGEN[2], the probability of a call being incorrect is reformulated to be more informative, however, the probabilities are formulated differently for homozygous reference, heterozygous, and homozygous alternative calls which give each a different distribution. In addition, the GQ is then rescaled during joint-calling, which will change the distribution depending on the method used (**Supplementary Figure 5**). For UKBB, DRAGEN was also used for joint-calling, whereas for AoUv8 the Genomic Variant Store[3] was used, followed by export to Hail VDS[4]. The distributions between the two cohorts were also affected for homozygous reference calls because AoUv8 pre-filtered homozygote reference genotypes with GQ<20 and rounded down the remaining values to the nearest 10 with a maximum of GQ=40.

##### *Other metrics of data quality*

Transitions (Ti) refer to base pair substitutions between a one-ring pyrimidine (C↔T) or between a two-ring purine (A↔G), and transversions (Tv) refer to base pair substitutions between a purine and a pyrimidine. The Ti/Tv ratio is often used for QC of sequencing data. For exome sequencing the observed Ti/Tv ratio is ~5, whereas for whole genome sequencing the expected value ~2.1[5]. The autosomal Ti/Tv ratio per individual after QC was similar across the QC methods, with the mean TiTv of 2.0758, 2.0706, 2.0805, 2.0783 for UKBB-GenotypeQC, UKBB-DRAGEN-ML, AoU-GenotypeQC, and AoU-VETS respectively (**Supplementary Figure 7**), an increase of 0.25% and 0.11% in UKBB and AoU respectively when applying genotype-level QC (**Supplementary Table 1**).

The expected number of singleton genotypes per individual is highly variable, depending on factors including the population structure within the cohort. Having a poor singleton non-ref genotype call will be the difference between observing the variant or not[6]. We found that allele-level QC and the genotype-level QC singleton calls generally agreed with one another (Pearson correlation $r = 0.9729$, $p < 10^{-308}$, and $r=0.9998$, $p<10^{-308}$ for UKBB and AoU respectively) (**Figure 2**). These differences were not driven by deviation from EUR-like genetic ancestry, which results in a higher overall singleton count (means by genetic ancestry groups of 1022.3 and 1013.1 (African, AoU), 5710.7 and 5388.5 (African, UKBB), 1158.1 and 1153.0 (Admixed American, AoU), 9497.1 and 9142.6 (Admixed American, UKBB), 4324.8 and 4328.7 (East Asian, AoU), 8823.3 and 8461.5 (East Asian, UKBB), 5205.6 and 5213.1 (Middle Eastern, AoU), 1207.1 and 1199.7 (European, AoU), 826.6 and 718.2 (European, UKBB), 5165.5 and 5173.8 (South Asian, AoU), and 4836.7 and 4607.6 (South Asian, UKBB) per individual for the allele-level and genotype-level QC sets respectively, **Supplementary Table 2**). Overall, 95%, and 63% of outliers (>5 standard deviations from the mean difference) were of European genetic ancestry in UKBB and AoUv8 respectively, which is a slight enrichment of outliers in genetically European individuals in AoUv8 (OR = 1.123 [1.035 - 1.219], p = 0.005), but not significantly different from the background distribution of genetic ancestries in UKBB (OR=0.996 [0.915 - 1.085], p=0.93).

#### *Summary of AoUv8 VDS release*

In AoUv8 the starting point for applying genotype-level QC is the provided Hail VDS[4,7]. This VDS consists of two Hail matrix tables, a 'reference_data' table, which stores blocks of hom-ref genotype calls with the same GQ value, and a 'variant_data' table, which stores non-hom-ref genotype calls with per-call metrics as well as per-site filter sets. The VETS filter[8] is represented by a boolean FT entry for each individual genotype call, but because VETS is actually an allele-level method FT is the same for all individuals with the same genotype at a given locus. In the case of multi-allelic variants where het-alt genotypes are possible, FT should be 'fail' if one or both of the alt alleles failed, although this rule was not always applied correctly in the AoUv8 release, see patch 0.6.3 notes in the Genomic Variant Store (GVS) changelog [https://github.com/broadinstitute/gatk/blob/ah_var_store/scripts/variantstore/docs/CHANGELOG.md].

We found that the most cost effective way of converting the VDS to genotype-QC'ed plink2 format[9], for use with downstream tools, was to export sharded VCFs from Hail (with FILTER, GT, LAD, GQ, FT fields) and then apply QC, convert and merge with our modified version of plink2. Exporting the VCFs from Hail cost ~£3500 and conversion with plink cost ~£350. We tested applying our QC filters in Hail before exporting, but we estimated that this would cost significantly more overall. We would not recommend exporting the VDS genome-wide to dense formats without a 'sharded' export option, such as plink and bgen, because this does not take advantage of the blocked nature of the VDS, and is slow and expensive.

The process by which hom-ref GQ values were blocked to {0, 20, 30, 40} and the GQ < 20 block set to missing is not explicitly documented in the AoU release notes[10]. However, the GVS github [https://github.com/broadinstitute/gatk/blob/ah_var_store/scripts/variantstore/] indicates that the v8 release gVCFs were reblocked and GQ < 20 hom-ref calls were dropped at the point of reblocked gVCF ingest into the GVS, as described in [https://github.com/broadinstitute/gatk/blob/ah_var_store/scripts/variantstore/docs/aou/AOU_DELIVERABLES.md] "NOTE Be sure to set the input drop_state to "ZERO" (this will have the effect of dropping GQ0 reference blocks)".

### Supplementary Figures

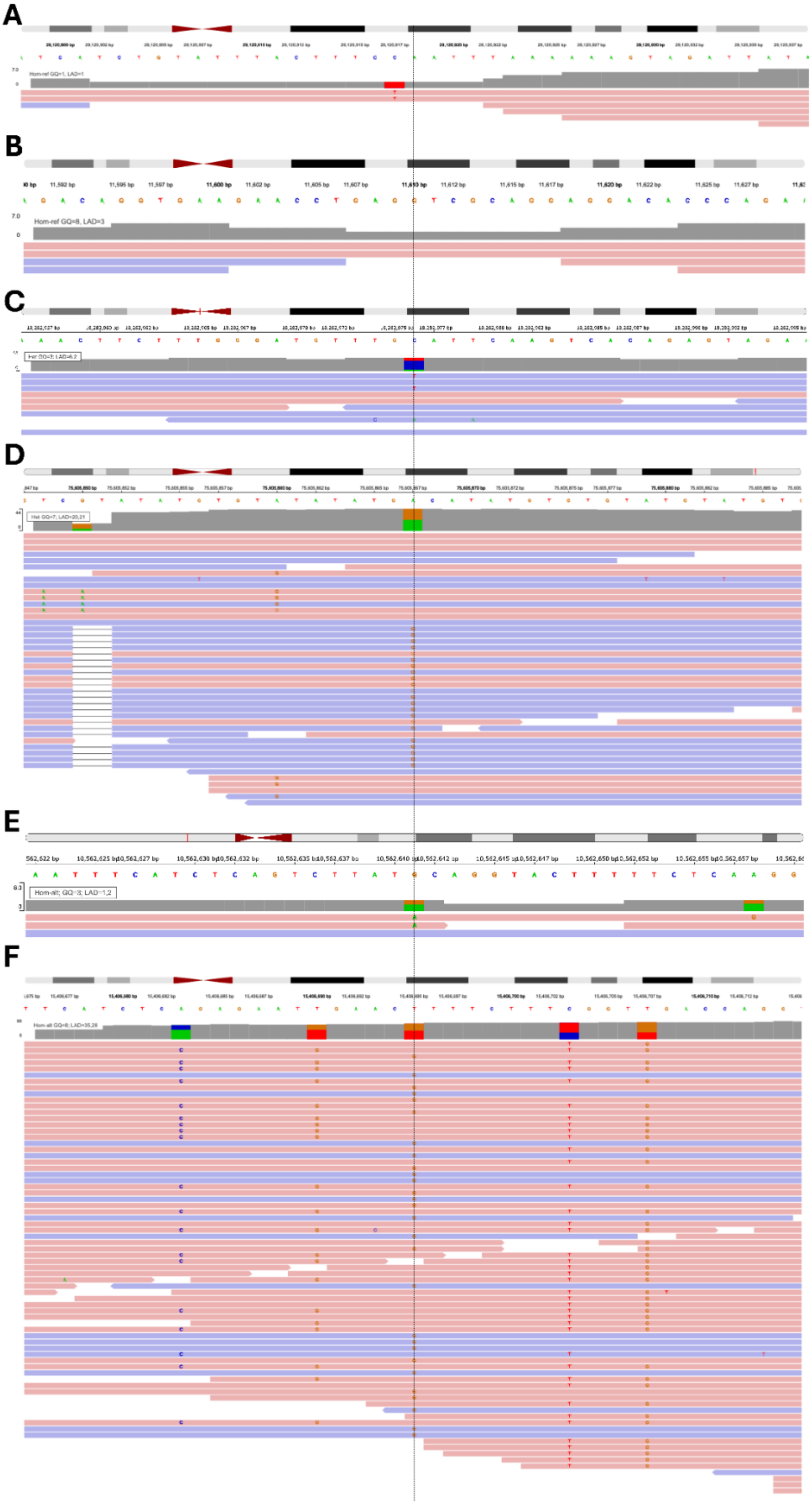

**Supplementary Figure 1: Read-level review of erroneous calls in UK Biobank** The dotted line shows the position of the calls. A) and B) are homozygous reference calls with a GQ=8 and 3 respectively, and an LAD=1 and 3 respectively. Of note, due to the blocking of hom-refs, the allelic depth and GQ is not for the individual call which can lead to different numbers of reads in LAD versus what is actually there. C) and D) are heterozygous genotypes with GQ=3 and 7, and LAD=6,2 and 20,21 respectively. The former has too few reads to be a confident call and has a second alt genotype called in one of the six 'reference' reads. The latter is an interesting genotype which is filtered out by our GQ filter, which is part of a complex haplotype which can be seen with the indel upstream appearing in the same reads, this is an example of a good call which we would remove as a consequence of filtering at scale. E) and F) are hom-alt calls with GQ=3 and 8, and LAD=1,2 and 35,28 respectively. The former has too few reads to make a variant call, but the latter appears to be a miscalled heterozygous genotype. Data visualised using IGV[11].

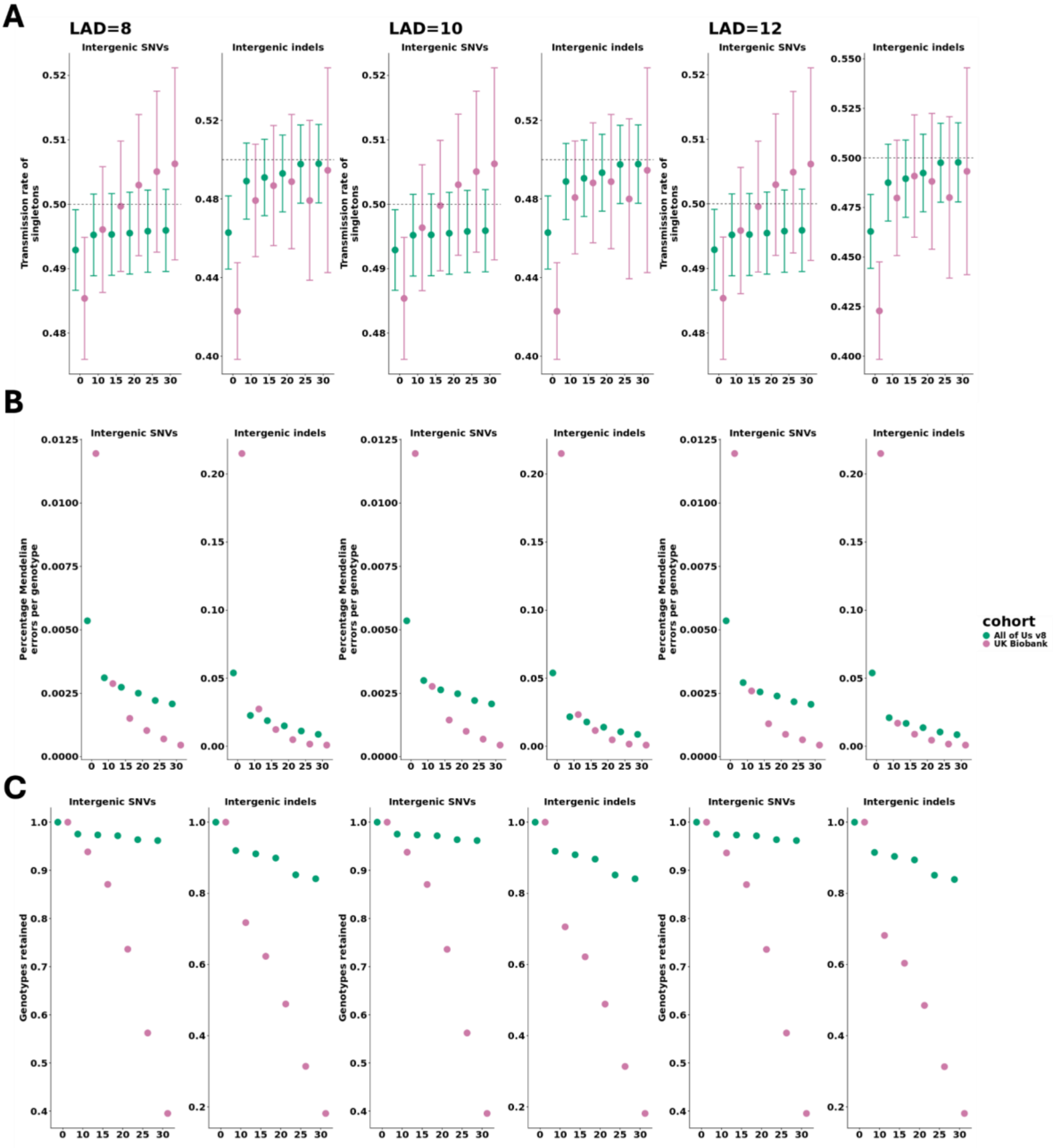


**Supplementary Figure 3: Transmission rate, Mendelian error rate, and genotypes retained for different genotype-level QC thresholds on 100 randomly selected autosomal pVCF chunks**

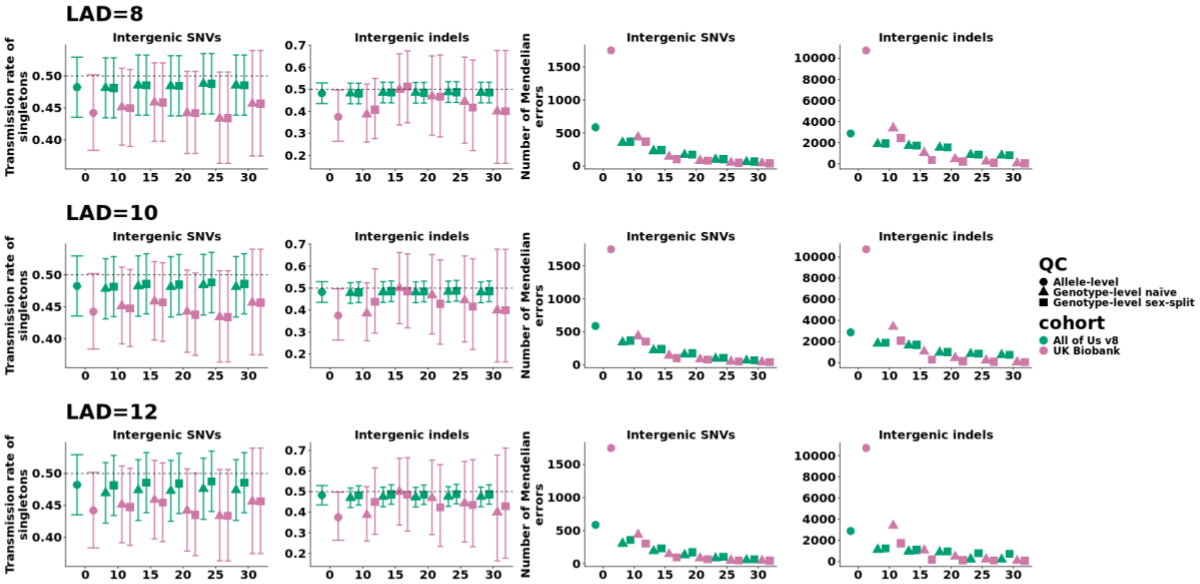


**Supplementary Figure 4: Transmission rate of singletons in male probands and the number of Mendelian errors using different QC thresholds on the X chromosome from 10 randomly selected vcf chunks on the non-pseudoautosomal regions**

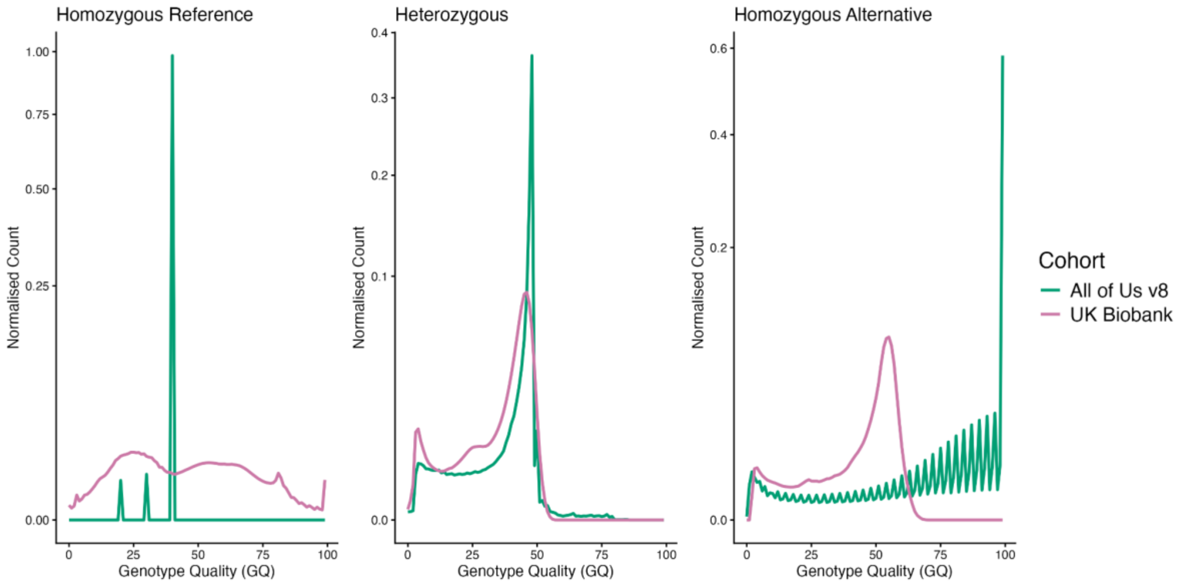


**Supplementary Figure 5: GQ distributions for homozygous reference, heterozygous, and homozygous alternative genotypes from a random sampling of variants across All of Us v8 and UK Biobank**

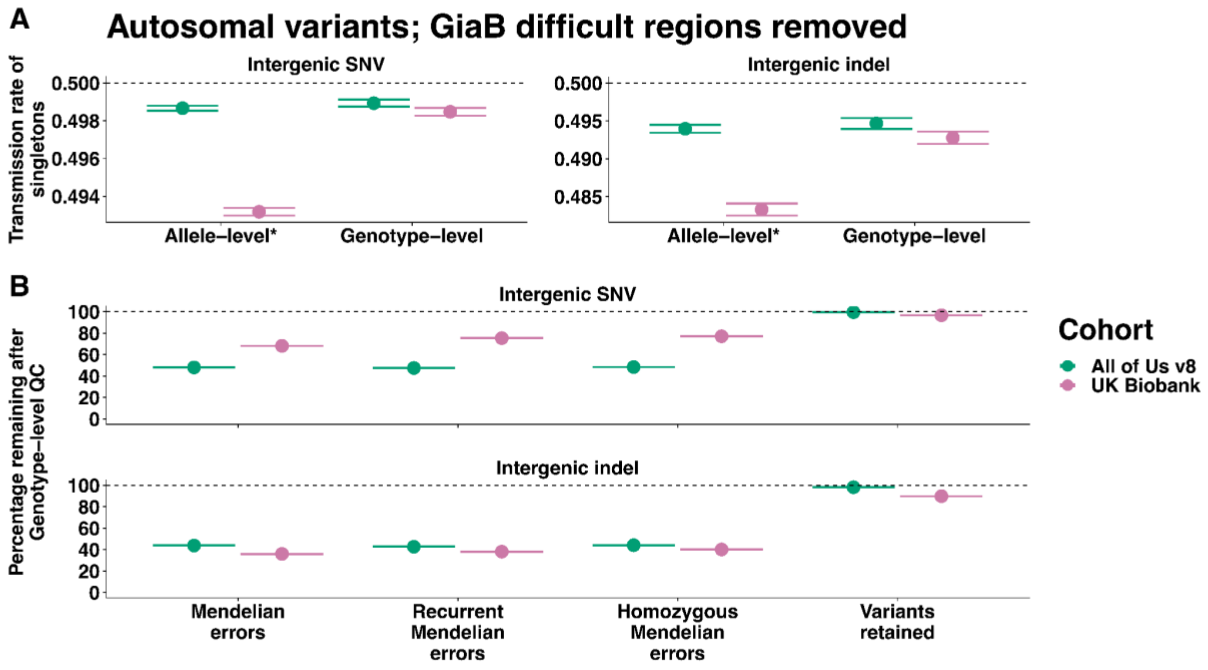


**Supplementary Figure 6: Transmission rate, percentage of errors remaining, and variants retained after genotype-level QC after removing difficult regions defined by Genome in a Bottle (GiaB)**

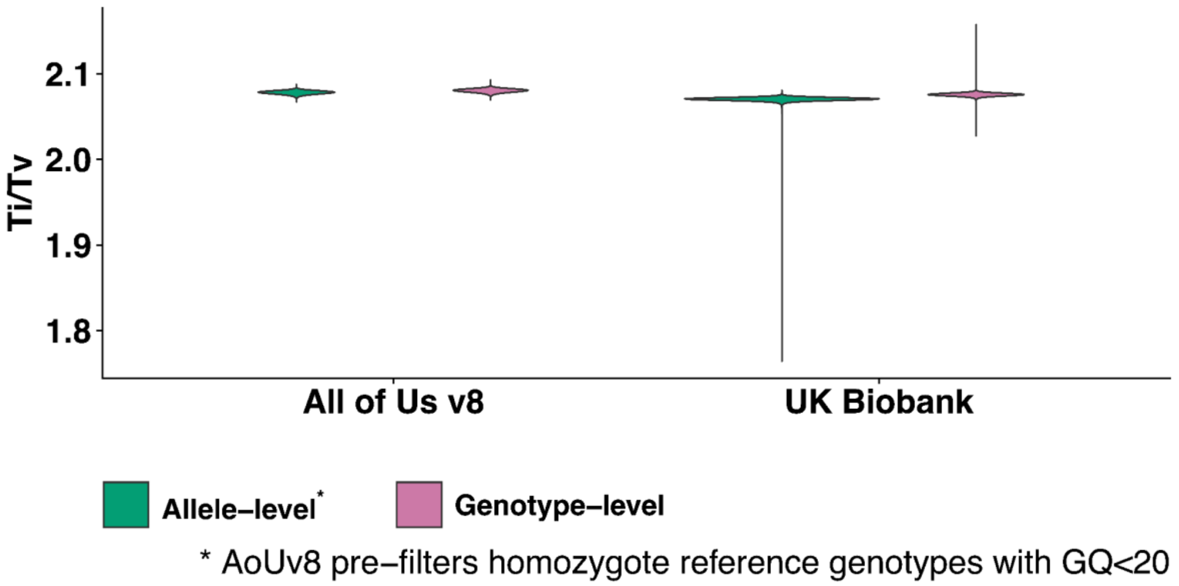


**Supplementary Figure 7: Transition to transversion distributions across the autosome for All of Us v8 and UK Biobank**

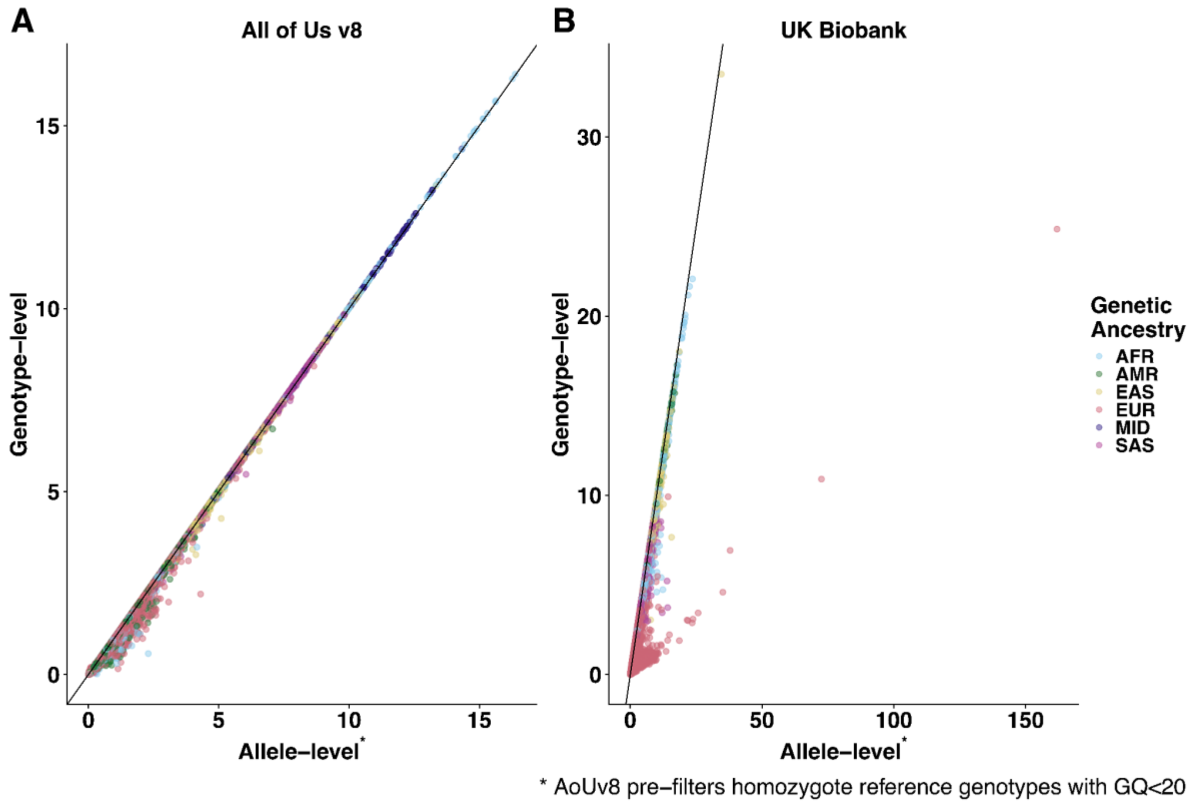


**Supplementary Figure 8: Number of singletons per individual across the autosome for All of Us v8 and UK Biobank from the Allele-level QC and Genotype-level QC datasets. The colour indicates the inferred genetic ancestry.**
